# Non-catalytic binding sites induce weaker long-range evolutionary rate gradients than catalytic sites in enzymes

**DOI:** 10.1101/554436

**Authors:** Avital Sharir-Ivry, Yu Xia

## Abstract

Enzymes exhibit a strong long-range evolutionary constraint that extends from their catalytic site and affects even distant sites, where site-specific evolutionary rate increases monotonically with distance. While protein-protein sites in enzymes was previously shown to induce only a weak conservation gradient, a comprehensive relationship between different types of functional sites in proteins and the magnitude of evolutionary rate gradients they induce has yet to be established. Here, we systematically calculate the evolutionary rate (dN/dS) of sites as a function of distance from different types of binding sites on enzymes and other proteins: catalytic sites, non-catalytic ligand binding sites, allosteric binding sites, and protein-protein interaction sites. We show that catalytic binding sites indeed induce significantly stronger evolutionary rate gradient than all other types of non-catalytic binding sites. In addition, catalytic sites in enzymes with no known allosteric function still induce strong long-range conservation gradients. Notably, the weak long-range conservation gradients induced by non-catalytic binding sites on enzymes is nearly identical in magnitude to those induced by ligand binding sites on non-enzymes. Finally, we show that structural determinants such as local solvent exposure of sites cannot explain the observed difference between catalytic and non-catalytic functional sites. Our results suggest that enzymes and non-enzymes share similar evolutionary constraints only when examined from the perspective of non-catalytic functional sites. Hence, the unique evolutionary rate gradient from catalytic sites in enzymes is likely driven by the optimization of catalysis rather than ligand binding and allosteric functions.

## Introduction

Enzymes are key players in metabolic pathways and are crucial for cell function. They bind their chemical reactants and enhance the production rate of chemical products by several orders of magnitude. The catalytic function occurs in the catalytic site, which is usually composed of relatively buried residues in a large cleft ^1^. Due to their function, catalytic site residues are highly conserved and even their close vicinity residues are under strong selective pressure where their conservation decreases with distance from the catalytic site ^1^. Recently it was shown that enzymes exhibit a long-range, nearly-linear conservation gradient from their catalytic site that extends even up to ~30Å in distance ^2^.

Possible confounding factors of the observed long-range conservation gradient in enzymes include various local structural properties known to drive evolutionary rate variation of protein sites ^3–12^. The main local structural determinants known are residue solvent accessibility and residue packing ^6,8,9^ such that residues are generally less conserved with increasing solvent exposure or decreasing packing. Indeed, catalytic sites are generally relatively buried and therefore could induce a conservation gradient based solely on solvent exposure gradients. However, for enzymes where the catalytic site is on the surface, a strong conservation gradient was observed as well ^2^, demonstrating that distance and solvent exposure contribute independently to the conservation gradient. Still, given that protein fold usage is very different between enzymes and non-enzymes ^13–15^, it is possible that other local structural determinants (other than solvent exposure and residue packing) could potentially cause the strong conservation gradient from catalytic sites in enzymes. However, we have previously shown that catalytically-inactive pseudoenzymes with nearly identical tertiary structure as enzymes exhibit a significantly reduced conservation gradient, thereby demonstrating that the observed conservation gradient in enzymes cannot be dominated by any backbone-based local structural determinants ^16^.

This study addresses the following outstanding question: Are catalytic sites unique in their abilities to induce such strong long-range conservation gradients? What about other non-catalytic functional sites which share similar functional capacities of binding another molecule as catalytic binding sites? Using off-lattice protein model, it was shown that the requirement to maintain a specific ligand-binding site gives rise to a conservation gradient from the ligand-binding site ^17,18^. For protein-protein interaction sites however, it was shown that interfacial sites induce significantly weaker conservation gradients than catalytic sites in enzymes ^2^. Here, we present a data-driven proteome-level study of the evolutionary rate (dN/dS) of residues as a function of their distance from different types of functional sites in proteins. We consider four types of functional sites: catalytic sites, non-catalytic ligand binding sites, allosteric binding sites, and protein-protein interaction sites. We show that all types of non-catalytic binding sites induce significantly weaker long-range evolutionary rate gradients than catalytic binding sites in enzymes. Surprisingly, non-catalytic ligand-binding sites on enzymes do not induce significant long-range evolutionary rate gradients. Instead, the weak evolutionary rate gradient induced from non-catalytic ligand-binding sites on enzymes resembles that from ligand-binding sites on non-enzymes. Moreover, we show that catalytic sites in enzymes with no known allosteric function still induce a strong long-range evolutionary rate gradient, indicating that allosteric function is not the main driving force of the observed evolutionary rate gradients from catalytic sites. Lastly, we show that solvent exposure gradients cannot explain the differences in magnitude between evolutionary rate gradients induced from catalytic and non-catalytic ligand-binding sites. Taken together, our results suggest that the observed evolutionary rate gradient from catalytic sites in enzymes is primarily driven by the optimization and maintenance of catalytic function rather than ligand-binding or allosteric function.

## Results

### Non-catalytic ligand-binding sites induce weaker evolutionary rate gradients than catalytic binding sites

To examine whether evolutionary rate gradients are also induced from non-catalytic ligand-binding sites, we first identified such functional binding sites within enzymes and non-enzymes in the yeast proteome. As a starting point, we used a dataset of the yeast proteome that contains 1744 structurally annotated yeast proteins (see Methods). We screened them to find yeast proteins for which the structural model contains a biologically significant ligand bound to it based on the BioLip database ^19^, excluding ions. We found 486 ligand-binding sites on 240 enzymes and 462 ligand-binding sites on 276 proteins which are not known to be enzymes (non-enzymes). Ligand-binding residues were identified from BioLip, and catalytic site residues in enzymes were identified using the Catalytic Site Atlas ^20^. The ligand-binding sites on enzymes were then subdivided into 250 ligand-binding sites that include catalytic site residues, and 236 ligand-binding sites that are completely separated from the catalytic site. On average, the evolutionary rates of the structurally modeled non-enzymes and enzymes in our datasets are not significantly different (0.0676±0.0009 and 0.0656 ±0.0007 respectively). For each residue, we calculated the distance to the closest ligand-binding residue. Average evolutionary rate (dN/dS) was then calculated for residues over distance bins to obtain the evolutionary rate gradients.

The slope of the linear fit for the evolutionary rate gradient from catalytic binding sites is significantly larger than the slope for the evolutionary rate gradient from non-catalytic ligand-binding sites (t-test, P<0.001, Table 1 and Figure 1). Surprisingly, the evolutionary rate gradient induced from non-catalytic ligand-binding sites on enzymes is weak and actually resembles the evolutionary rate gradient induced from ligand-binding sites on non-enzymes (t-test, P=0.17). The weak evolutionary rate gradient from non-catalytic ligand-binding sites suggests that ligand-binding functionality of catalytic sites is probably not the main determinant of the strong evolutionary rate gradients induced from them.

**Table 1.** Non-catalytic ligand binding sites induce weaker evolutionary rate gradients than catalytic binding sites. Slope of the linear fit of the average evolutionary rate (dN/dS) versus distance from catalytic sites and non-catalytic ligand-binding sites, as well as the average evolutionary rate of binding site residues.

|  | Type of binding site | Slope | Evolutionary rate (dN/dS) of binding site residues |
| --- | --- | --- | --- |
| Enzymes | Catalytic ligand-binding sites | 0.0035<br>(±0.0003) | 0.011 (±0.002) |
| Enzymes | Non-catalytic ligand-binding site | 0.0018<br>(±0.0001) | 0.030 (±0.004) |
| Non-enzymes | Non-catalytic ligand-binding site | 0.0015<br>(±0.0002) | 0.0312 (±0.003) |

- In brackets – standard errors

**Figure 1.**
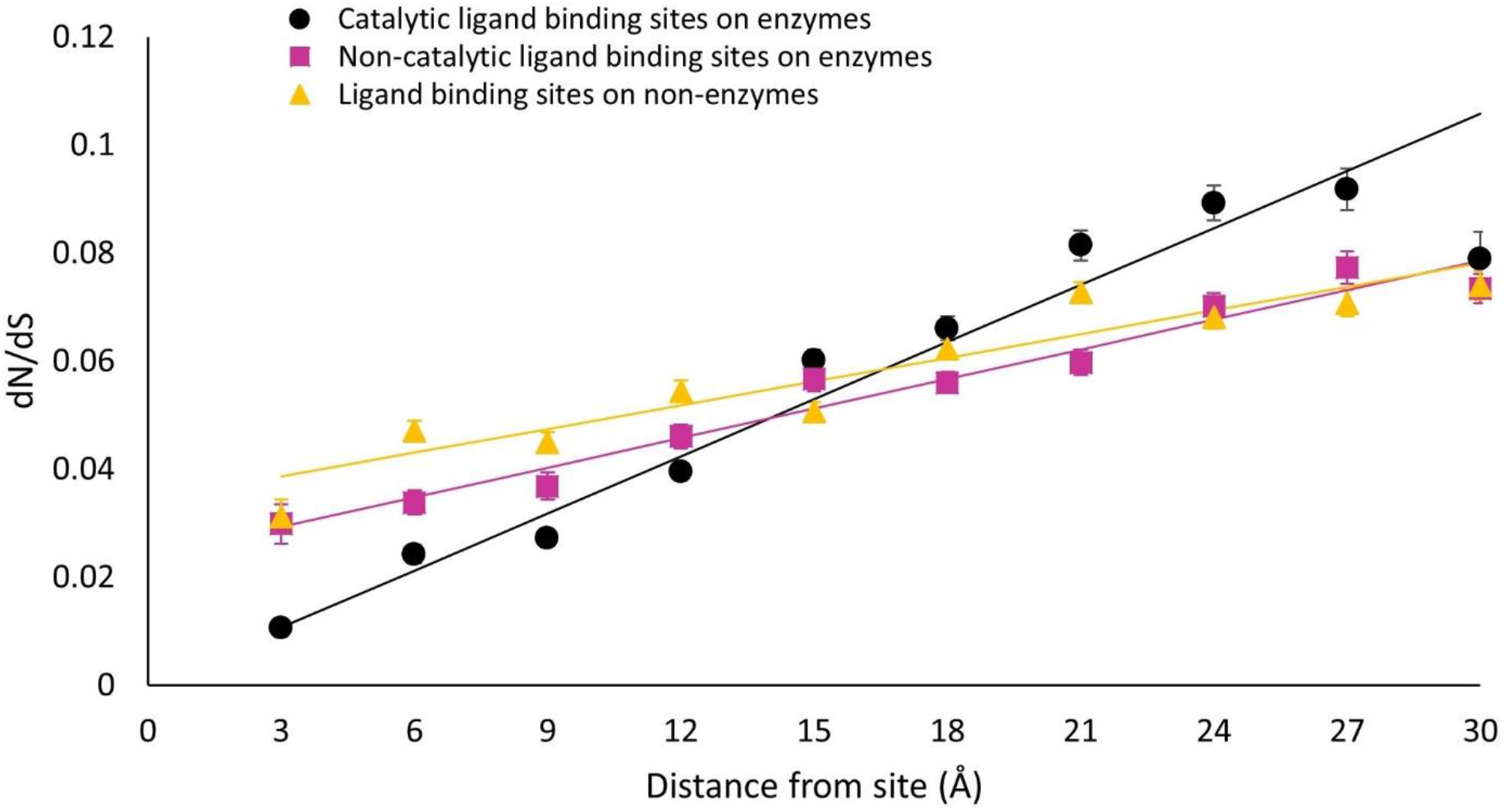
Non-catalytic ligand binding sites induce weaker evolutionary rate gradients than catalytic binding sites. Evolutionary rate (dN/dS) as a function of distance from catalytic and non-catalytic ligand-binding sites in enzymes, as well as from ligand-binding sites in non-enzymatic proteins.

### Allosteric binding sites induce weaker long-range evolutionary rate gradients than catalytic sites

Next, we focused on the special case of allosteric binding sites. We collected those yeast proteins that have a structural model which is known to have an allosteric function with its allosteric site residues annotated in the allosteric database (ASD) ^21^. We found 190 allosteric proteins, of which 108 are known enzymes with known catalytic site residues. All residues were binned according to their distance from the closest allosteric binding residue as well as from the closest catalytic residue in the case of enzymes. Average evolutionary rate (dN/dS) was calculated for the residues in each distance bin.

The slope of the evolutionary rate gradient induced from allosteric binding sites is significantly smaller than that induced from catalytic sites (t-test, P<0.001, Figure 2 and Table 2). Allosteric binding sites in enzymes modulate the activity of catalytic sites in that the catalytic site is shifting into its active conformation upon a binding event in the distant allosteric binding site ^22,23^. Despite this long-range interaction between catalytic sites and allosteric binding sites, our results suggest that allosteric function is not the main determinant of the long-range evolutionary rate gradient induced from catalytic sites in enzymes.

**Figure 2.**
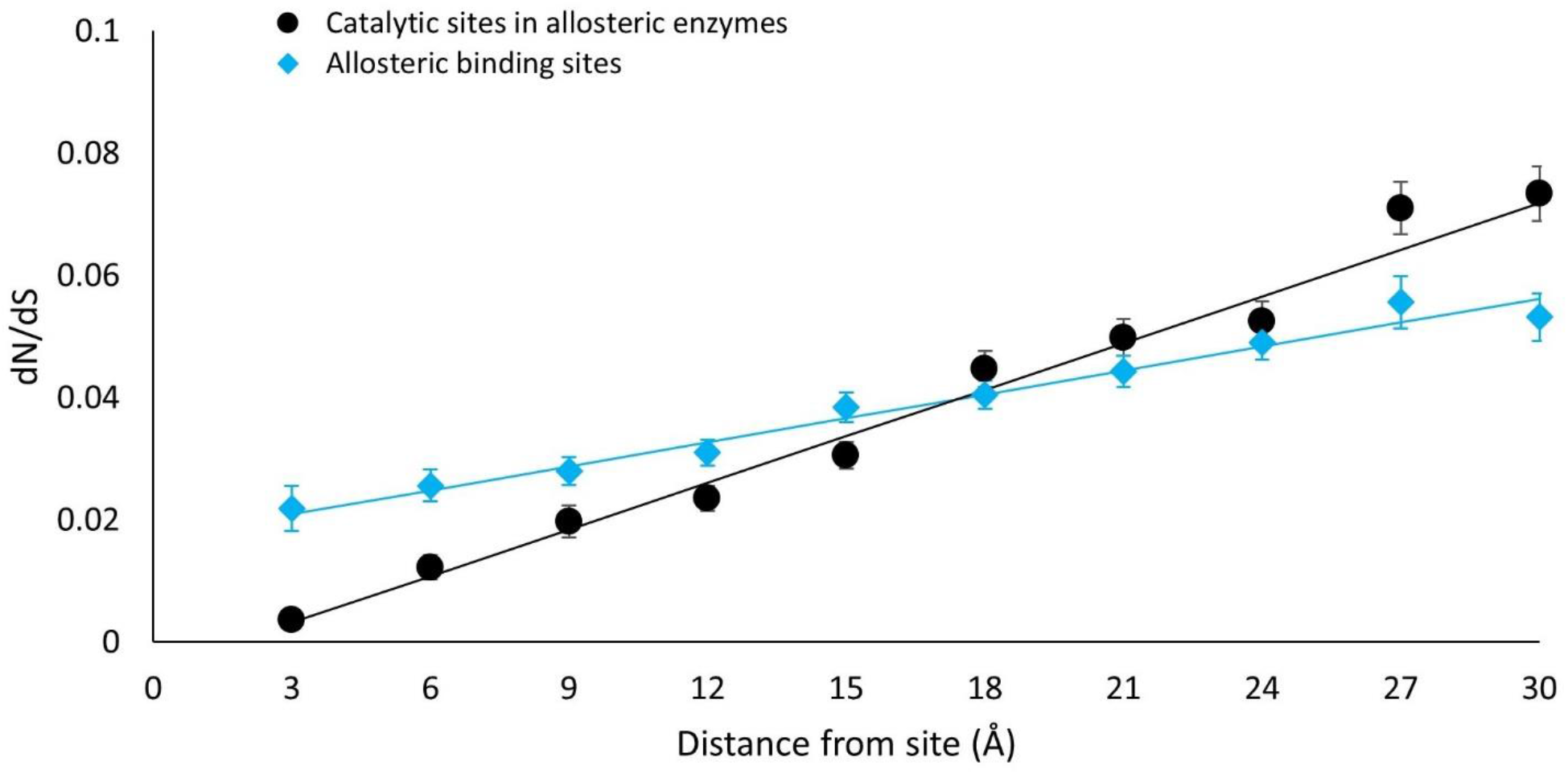
Allosteric binding sites induce weaker evolutionary rate gradients than catalytic sites. Evolutionary rate (dN/dS) as a function of distance from allosteric binding sites and from catalytic sites in allosteric enzymes.

**Table 2.** Allosteric binding sites induce weaker evolutionary rate gradients than catalytic sites and catalytic sites in non-allosteric enzymes induce strong evolutionary rate gradients. Slope of the linear fit of the average evolutionary rate (dN/dS) versus distance from allosteric binding sites and catalytic sites in allosteric enzymes, as well as the average evolutionary rate of binding site residues.

|  | Type of functional site | Slope | Evolutionary rate (dN/dS) of functional site residues |
| --- | --- | --- | --- |
| Allosteric enzymes | Catalytic site | 0.0025<br>($\pm 0.0001$) | 0.004 ($\pm 0.002$) |
| Allosteric enzymes | Allosteric binding sites | 0.0013<br>($< 0.0001$) | 0.022 ($\pm 0.004$) |
| Non-allosteric enzymes | Catalytic site | 0.0038<br>($\pm 0.0002$) | 0.010 ($\pm 0.004$) |

- In brackets – standard errors

### Yeast-annotated non-catalytic ligand-binding sites induce weaker evolutionary rate gradients than catalytic binding sites

The annotations of functional binding sites were made according to the homology-based structural models of the *Saccharomyces cerevisiae* proteins. These models are usually based on solved structures of homologous proteins in other species. In order to rule out the confounding factor that some of our homology-based annotations of functional sites are false positives in yeast, we examined a subset of yeast proteins which are also known to have the relevant binding functionality in yeast. The results for these yeast proteins with high-confidence annotations in Figure 3 and Table 3 clearly show a significant difference between the strong evolutionary rate gradients induced from catalytic binding sites on enzymes and all other non-catalytic binding sites. Due to the significantly smaller size of the yeast-annotated allosteric site dataset (only 11 sites), we reduced the number of distance bins to five and the slope is compared to the slope of the gradient induced from catalytic binding sites based on five bins as well. Overall, our results are robust to possible false positives of our homology-based functional binding site annotations for yeast proteins.

**Figure 3.**
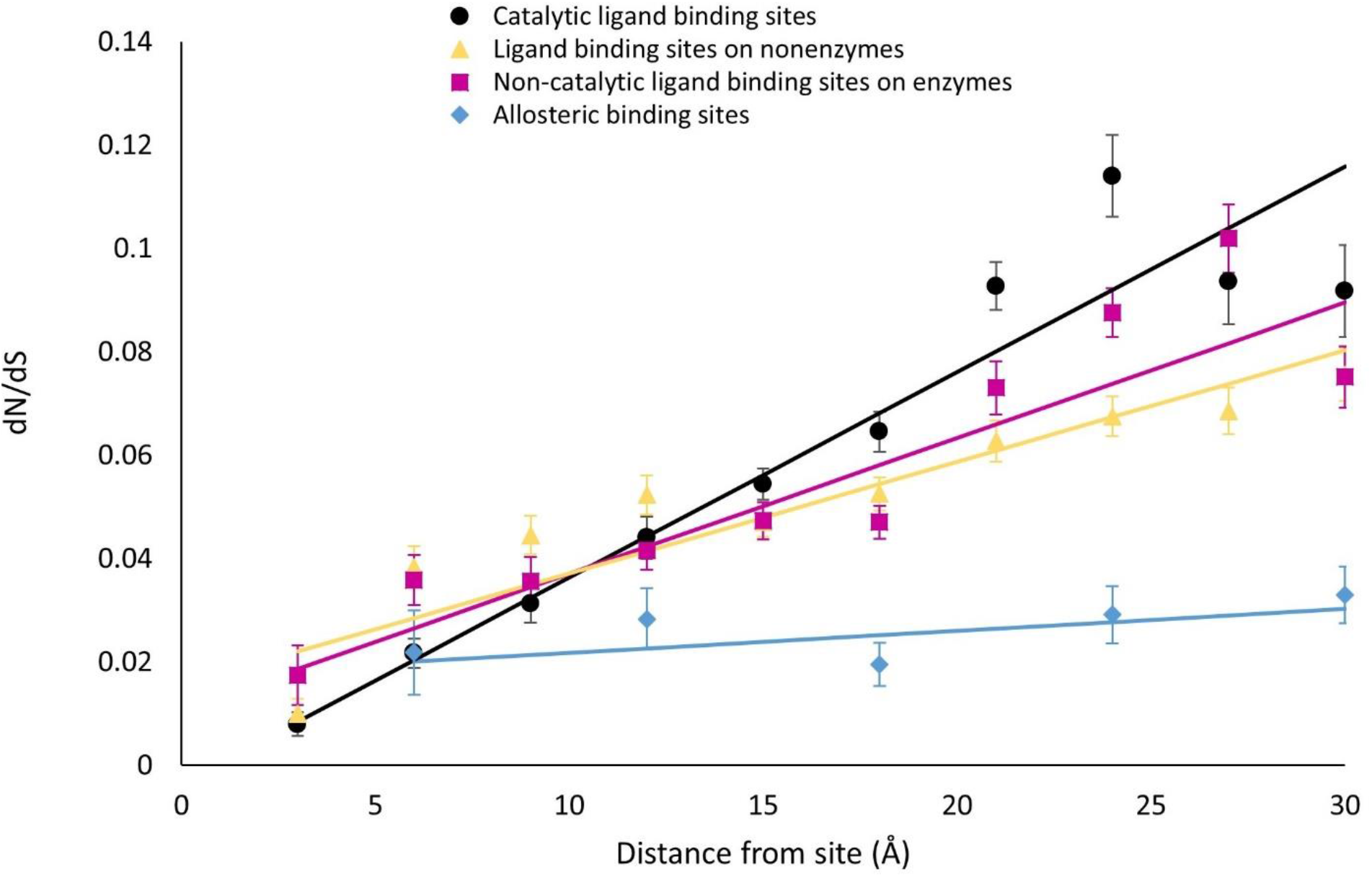
Yeast-annotated non-catalytic ligand binding sites induce weaker evolutionary rate gradients than yeast-annotated catalytic binding sites. Evolutionary rate (dN/dS) as a function of distance from catalytic and non-catalytic ligand-binding sites in enzymes as well as from ligand-binding sites in non-enzymatic proteins and allosteric binding sites for the subset of *Saccharomyces cerevisiae* proteins known to contain the respective functional binding sites.

**Table 3.** Yeast-annotated non-catalytic ligand binding sites induce weaker evolutionary rate gradients than yeast-annotated catalytic binding sites. Slope of the linear fit of the average evolutionary rate (dN/dS) versus distance from catalytic sites and non-catalytic ligand-binding sites, as well as the average evolutionary rate of binding site residues for yeast proteins known to contain functional binding sites.

| Type of binding site | Slope | Evolutionary rate (dN/dS) of binding site residues |
| --- | --- | --- |
| Catalytic binding site | 0.0040 ($\pm 0.0003$)<br>0.0037($\pm 0.0005$)<sup>#</sup> | 0.008 ($\pm 0.002$) |
| Non-catalytic ligand binding sites on enzymes | 0.0026 ($\pm 0.0005$) | 0.017 ($\pm 0.006$) |
| Ligand binding sites on non-enzymes | 0.0022 ($\pm 0.0003$) | 0.010 ($\pm 0.003$) |
| Allosteric binding sites | 0.0004 ($\pm 0.0003$)<sup>#</sup> | 0.022 ($\pm 0.008$) |

• In brackets – standard errors
<sup>#</sup> Based on slope calculated over 5 distance bins of 6 Å

### Catalytic sites in non-allosteric enzymes induce strong evolutionary rate gradients

For comparison, we also identified ‘non-allosteric enzymes’ in our dataset as those proteins with an enzyme structural model with known catalytic residues but with no known allosteric function (219 proteins). Notably, similar to allosteric enzymes, catalytic sites in non-allosteric enzymes also exhibit a strong evolutionary rate gradient that extends to distant sites (Figure 4 and Table 2). The existence of evolutionary rate gradient from catalytic sites appears to be independent of the existence of allosteric function in the enzyme. This result further supports the hypothesis that the allosteric function is not the main determinant of evolutionary rate gradients induced from catalytic site in enzymes.

**Figure 4.**
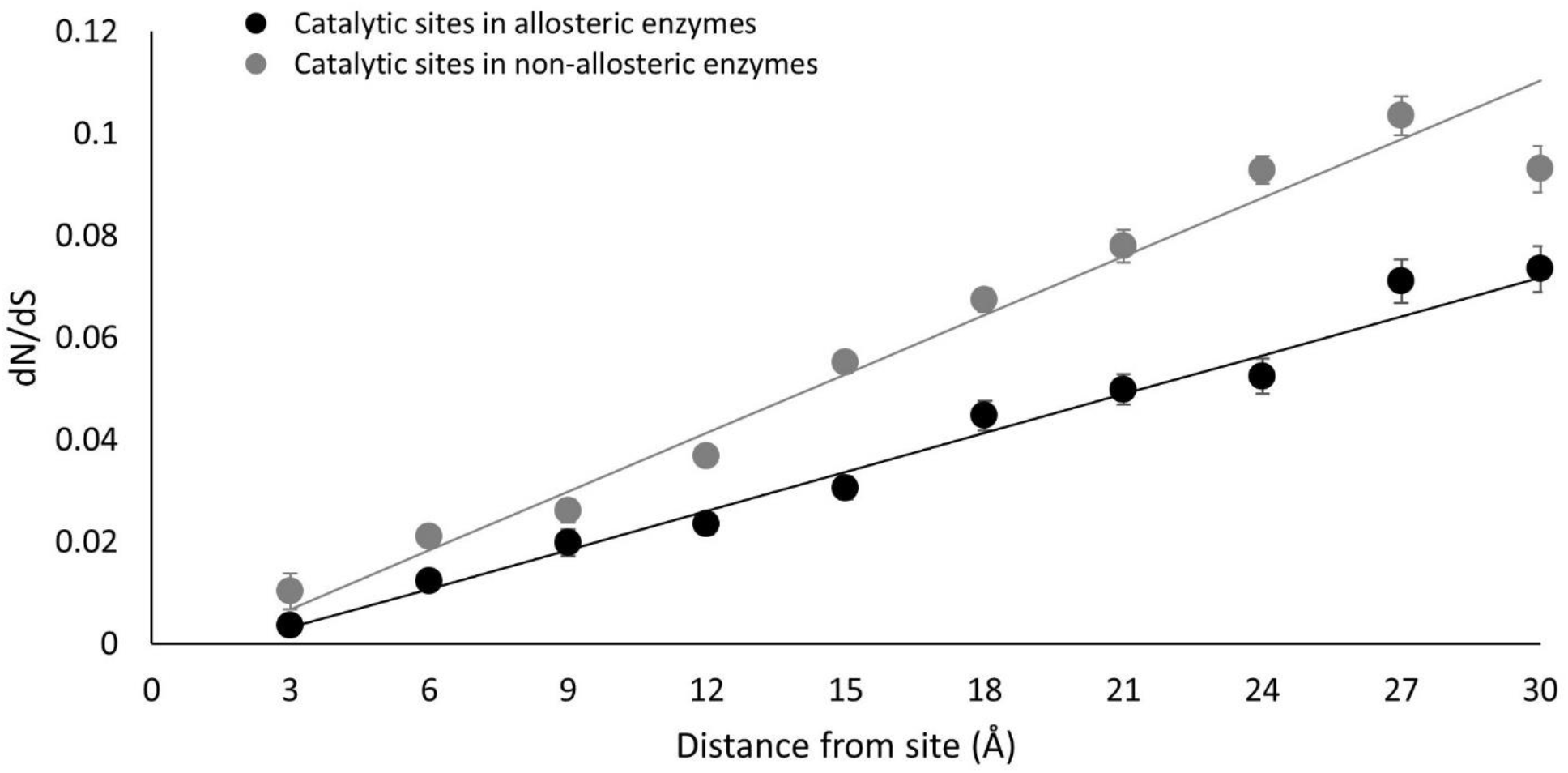
Catalytic sites in non-allosteric enzymes induce strong evolutionary rate gradients. Evolutionary rate (dN/dS) as a function of distance from catalytic sites in allosteric and non-allosteric enzymes.

### Protein-protein interaction sites induce weaker long-range evolutionary rate gradients than catalytic binding sites

Using relative conservation scores, it was previously shown that protein-protein interactions sites on enzymes induce only a minor conservation gradient which is significantly weaker than that from catalytic sites ^2^. Here, we studied residue evolutionary rate (dN/dS) as a function of distance from protein-protein interaction sites in the yeast proteome. We screened the structurally-annotated yeast proteome dataset to find proteins that are known to interact with other proteins (see Methods). We found 459 interfacial sites for 250 proteins. Interfacial residues were identified as those residues that that have different solvent accessibility when in complex compared to when the interacting partner is deleted. Residues were binned according to their distance from interfacial residues, and average dN/dS was calculated for each bin. The slope of the evolutionary rate gradient induced from interfacial sites is indeed significantly smaller than that induced from catalytic binding sites (t-test, P<0.001, Figure 5 and Table 4), either on enzymes or on non-enzymes.

**Figure 5.**
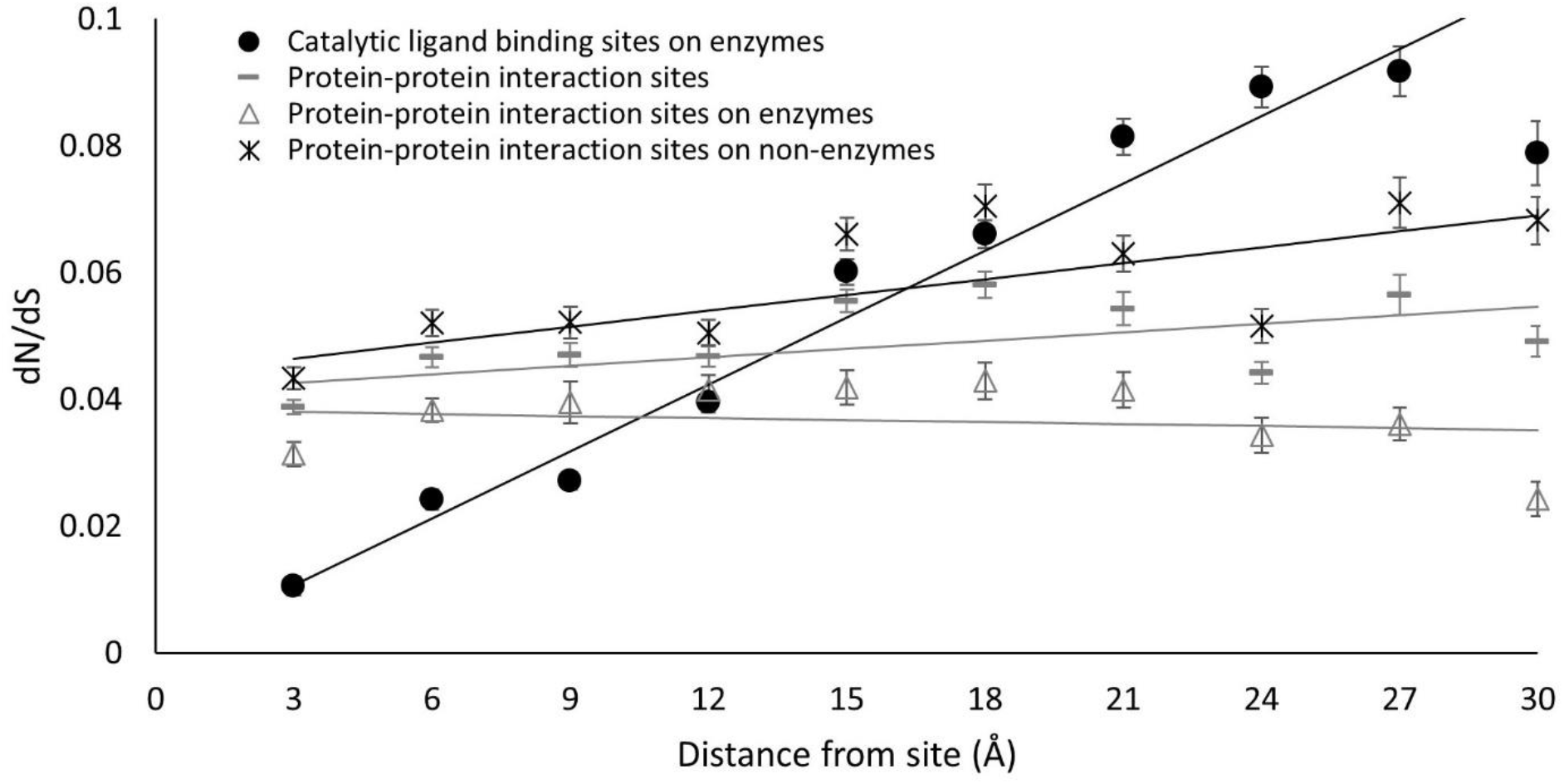
Protein-protein interaction sites on enzymes or non-enzymes induce weaker long-range evolutionary rate gradients than catalytic binding sites. Evolutionary rate (dN/dS) as a function of distance from catalytic binding sites and from protein-protein interaction sites on enzymes and non-enzymes.

**Table 4.**
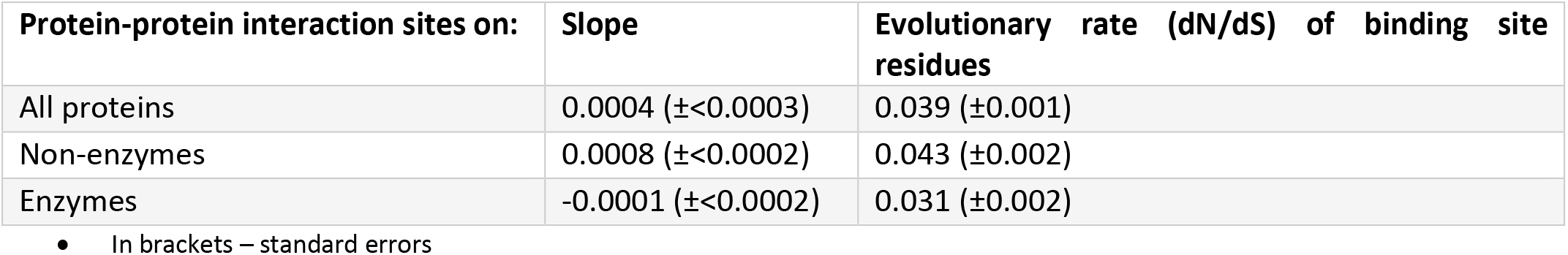
Protein-protein interaction sites on enzymes or non-enzymes induce weaker long-range evolutionary rate gradients than catalytic binding sites. Slope of the linear fit of the average evolutionary rate (dN/dS) versus distance from protein-protein interaction sites, as well as the average evolutionary rate of interfacial residues.

| Protein-protein interaction sites on: | Slope | Evolutionary rate (dN/dS) of binding site residues |
| --- | --- | --- |
| All proteins | 0.0004 ($\pm < 0.0003$) | 0.039 ($\pm 0.001$) |
| Non-enzymes | 0.0008 ($\pm < 0.0002$) | 0.043 ($\pm 0.002$) |
| Enzymes | -0.0001 ($\pm < 0.0002$) | 0.031 ($\pm 0.002$) |

- In brackets – standard errors

### Evolutionary rate gradient variations among different functional sites cannot be explained by local structural determinants

We examined whether local structural determinants such as solvent exposure gradients are responsible for the observed difference in magnitude of evolutionary rate gradients induced from catalytic and non-catalytic binding sites. We calculated the relative solvent accessibility (RSA) of each residue and then the expected linear trend of dN/dS versus RSA for each protein dataset. We then used the calculated slope and intercept of this dN/dS versus RSA relationship (fig. S1 and table S1 in the Supplementary Material) to plot the expected evolutionary rate at each distance bin according to the average RSA of the residues at that bin. As shown in Figure 6, there is no significant expected increase of evolutionary rate with distance based on solvent exposure alone for any of the datasets of functional binding sites. We therefore conclude that solvent exposure patterns cannot explain the differences in evolutionary rate gradients induced from catalytic and non-catalytic binding sites.

**Figure 6.**
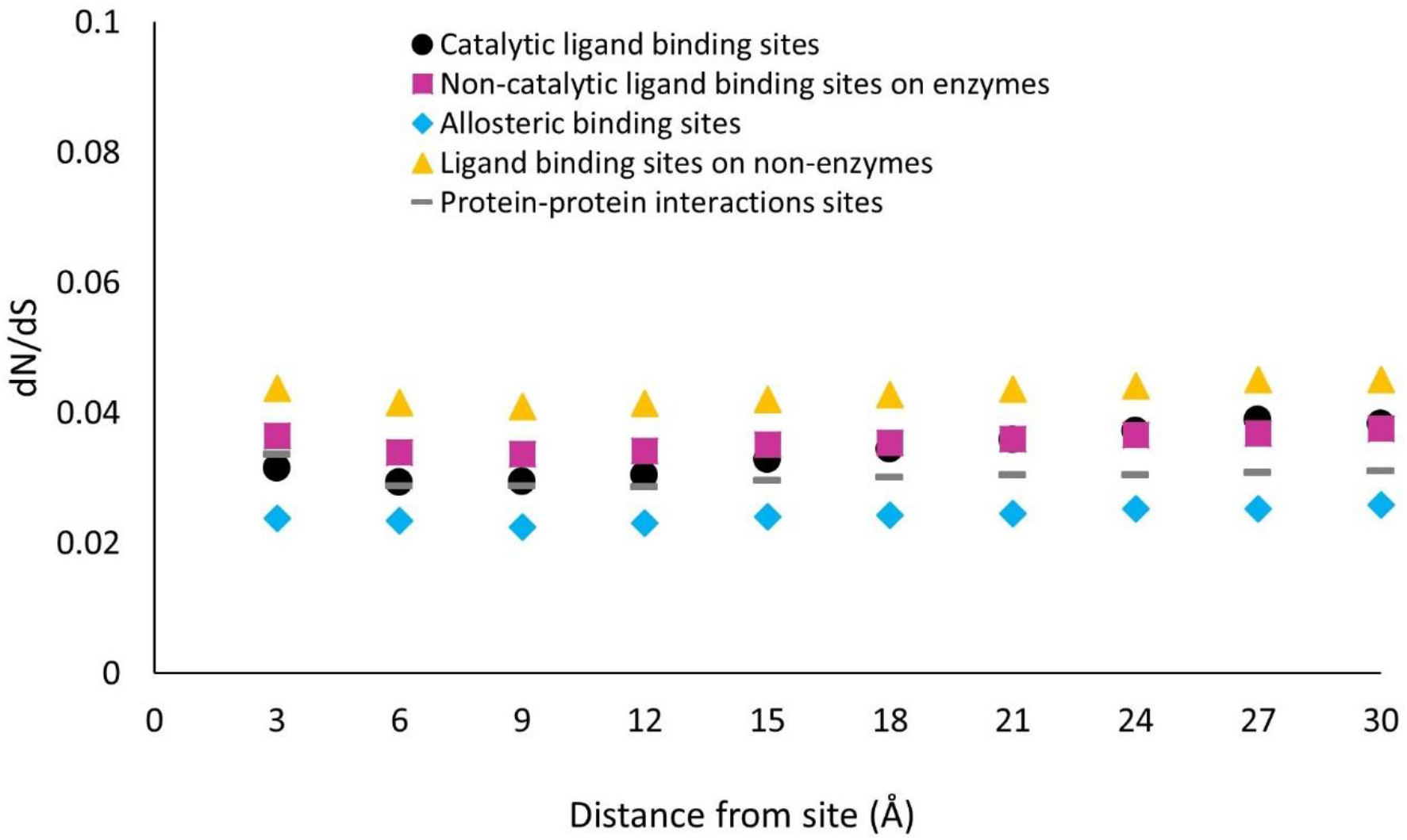
Evolutionary rate based on solvent exposure alone does not show an increase with distance from catalytic and non-catalytic binding sites. Expected dN/dS according to average relative solvent accessibility (RSA) of residues as a function of distance from binding site.

## Discussion

Strong long-range rate gradient is observed from catalytic sites in enzymes where there is a monotonic, nearly-linear increase in selective pressure on residues with their proximity to the catalytic site. We studied here functional non-catalytic ligand-binding sites on enzymes and non-enzymes to examine whether they induce similar evolutionary rate gradients as enzymatic catalytic sites. For this purpose, we used a structurally-annotated yeast proteome as a model to systematically study site-specific evolutionary rates as a function of distance from functional binding sites. We show that non-catalytic binding sites, even on enzymes, induce significantly weaker evolutionary rate gradients than catalytic sites.

Previous studies show that evolutionary rate gradients in enzymes cannot be explained by local structural properties of the protein ^2,16^, suggesting that the gradient is driven by functional determinants. A recent biophysical model of enzyme evolution that incorporates enzymatic activity constraint along with stability constraints better explains the evolutionary rate gradient from catalytic sites ^24^. Enzymatic function is very complex, involving both substrate binding and accelerating the production rate of chemical products by decreasing the free energy barrier of the chemical reaction. By showing that non-catalytic ligand-binding sites induce significantly weaker evolutionary rate gradients than catalytic sites, we can rule out several hypotheses on the origin of the long-range evolutionary rate gradient. The first hypothesis that can be ruled out is that the ligand-binding function of the catalytic site is the main determinant of the long-range rate gradient. While ligand binding is part of the function of the catalytic site, it appears improbable that the specific binding of the substrate is the main determinant of the long-range selection pressure in enzymes.

The second hypothesis that can be ruled out is that the long-range evolutionary rate gradient from catalytic sites is mainly driven by the allosteric function of the enzyme. Allosteric function exhibit a long-range effect in which binding of an effector molecule at a distant allosteric site shifts the catalytic site into an active conformation ^22,23^. It is reasonable to expect that the need to maintain proper long-range allosteric function can explain the observed long-range evolutionary rate gradient from the catalytic site. On the contrary, we have shown that allosteric binding sites induce a significantly weaker evolutionary rate gradient than catalytic sites. Moreover, a strong evolutionary rate gradient exists even in enzymes which are not known to be allosteric. These results suggest that optimizing the allosteric function of the enzyme is not the main determinant of the observed evolutionary rate gradient.

Finally, we show that for all the functional sites we have studied here, there is no significant solvent exposure gradients extending from them. Hence, the observed evolutionary rate gradient variations among different functional sites cannot be explained by known local structural determinants such as solvent exposure gradients.

We are therefore left with two plausible hypotheses on the origin of the unique evolutionary rate gradient from catalytic sites. The first hypothesis is that strongly conserved sites in general induce stronger long-range evolutionary rate gradient. Here, the evolutionary rate gradient is a result of a percolating effect of penetrating conservation ^18,25,26^. Thus, the higher the selective pressure on a site is, the stronger the percolation of evolutionary rates will be from it. The second hypothesis is that evolutionary rate gradients are driven by the chemistry of the enzyme where the main determinant of the increasing selective pressure is to optimize the catalytic power of the enzyme by specifically stabilizing the reaction transition state ^27^. Indeed, a chemistry-centered view on the evolution of new catalytic functions in enzymes suggests that new function in an enzyme can evolve from the ability of the enzyme to stabilize the transition state of the new chemical reaction (rather than just binding of the new ligand) ^28–30^. These two hypotheses are both consistent with the observations in this study, and they are not mutually exclusive. Further work is needed to critically examine the validity and applicability of these hypotheses in terms of explaining the origin of the long-range evolutionary rate gradient observed in enzymes.

## Methods

### Structural annotation of yeast proteins

We based our study on a dataset of structural homologs for the yeast proteome. This dataset was created using gapped BLAST ^31^ searches between protein subunit sequences with solved structure from the Protein Data Bank ^32^ and 5,861 translated open reading frames (ORFs) of the yeast *Saccharomyces cerevisiae* ^33^. We kept those ORF–subunit pairs in which both the subunit sequence and the ORF sequence had coverage of ≥50% in the alignment and E-value <10^-5^ and could be paired with their orthologs in three other closely related yeast species *S. paradoxus, S. mikatae*, and *S. Bayanus*. The subunit that produced the lowest alignment E-value with the ORF was chosen as the structural homolog. This way, 1798 yeast ORFs were mapped to a homolog in the PDB. The site-specific structural features of the PDB homolog we

#### Annotation of ligand-binding sites and catalytic sites

The structural subunits were screened to find those with an identified biologically-relevant ligand-binding site according to the BioLip protein function database ^19^. We excluded binding sites where the ligand is an ion. Yeast ORF was classified as an enzyme if its paired structural subunit has a known EC number. Catalytic sites of protein models were identified using the Catalytic Site Atlas ^20^. 240 enzymes with 486 ligand-binding sites were identified. We then subdivided the ligand-binding sites on yeast enzymatic proteins into those with an overlap between binding site and catalytic site residues (250 catalytic ligand-binding sites) and those where none of the binding site residues is a catalytic residue (236 non-catalytic ligand-binding sites). 276 proteins with 462 ligand-binding sites are not assigned with an EC number and were therefore termed as ‘non-enzymes’. A subset of the dataset which contains only yeast proteins which are known to contain the relevant functional binding sites was created as well and included 59 enzymes with 60 catalytic ligand binding sites and 75 non-catalytic ligand binding sites and 76 non-enzymes with 125 ligand binding sites. The distance of the Cα atom of each residue in these proteins to the closest Cα of the binding site residues and catalytic site residues was calculated. Residues were binned according these distances up until 30Å into equally spaced bins of 3Å in size. The first bin essentially contained all the ligand-binding residues and only them.

#### Annotation of allosteric proteins

Using the dataset of proteins with known allosteric function from the allosteric database (ASD) ^21^, we screened the yeast ORFs by means of their structural subunit to those that are known to have an allosteric function and their allosteric sites residues are known. 196 yeast ORFs matched this criterion, and out of them we kept 190 proteins for which the structural alignment contained the allosteric binding sites. Within the dataset, 108 proteins are identified as enzymes with an EC number and with known catalytic site from the Catalytic Site Atlas (Furnham et al., 2014). Other proteins in our dataset which have an EC number assigned and their catalytic site residues are identified, but whose structural models are not known to have an allosteric function (219 proteins), were classified as ‘non-allosteric’ enzymes. In addition we created which are known to have allosteric function in ASD. The distance of the Cα atom of each residue to the closest Cα atom of allosteric site residue and closest Cα atom of catalytic site residue was calculated. Residues were binned according these distances up until 30Å into equally spaced bins of 3Å in size and for the subset of 11 proteins into equally spaced bins of 6 Å to have sufficient number of residues in each bin.

#### Protein interfaces

We screened the protein complexes from which the best ORF-subunit were derived to those that are in physical contact with another ORF-subunit pair (not necessarily the pair with highest similarity) and were reported as interacting with the other ORF-subunit pair by at least one physical experiment in the BioGRID ^34,35^. Our dataset contained 250 yeast proteins with 459 interfaces. Interfacial residues were identified as the residues which have different solvent accessibility values when in complex compared to when the interaction partner is manually deleted.

All of the optimal ORF-subunit pairs and the corresponding subunit binding site residues are listed in supplementary table S2 in the Supplementary Material and their alignments in supplementary file S1.

### dN/dS calculations

We used previous alignments made between the translated ORFs from *S. cerevisiae* and their orthologs in three other closely related species *S. paradoxus, S. mikatae*, and *S. Bayanus* ^9^. The aligned codons were binned according to their distance from different binding sites as explained above and then concatenated. dN/dS values were calculated over the multiple sequence alignments using the program codeml within the PAML software package ^36^. As our four yeast species are closely related, we opted to calculate a single dN/dS value for the entire tree (using model=0), which we specified as ((*S. cerevisiae, S. paradoxus), S. mikatae, S. bayanus*). Codon frequencies were assumed equal (CodonFreq=0) and other parameters in codeml were left to their default values. The codon alignments can be found in supplementary files S2 in the Supplementary Material.

### Statistical analysis

For each bin, we estimated the standard error in our measurements of dN/dS using 50 rounds of bootstrap resampling. We used a weighted least squares regression to fit dN/dS versus distance where the standard errors of dN/dS in each bin was considered such that distance bins with small dN/dS estimation errors receive greater weight in the line fitting process. We used two-tailed t-tests to compute the significance of the difference between slopes.

## Acknowledgments

This work was supported by Natural Sciences and Engineering Research Council of Canada (grant number RGPIN-2014-03892 to Y.X.), Canada Research Chairs program (to Y.X.), Canada Foundation for Innovation (grant numbers JELF-33732, IF-33122 to Y.X.) and Dalia and Dan Maydan Post-doctoral Fellowship (to A.S.-I.).

## Supplementary material

**Figure S1.**
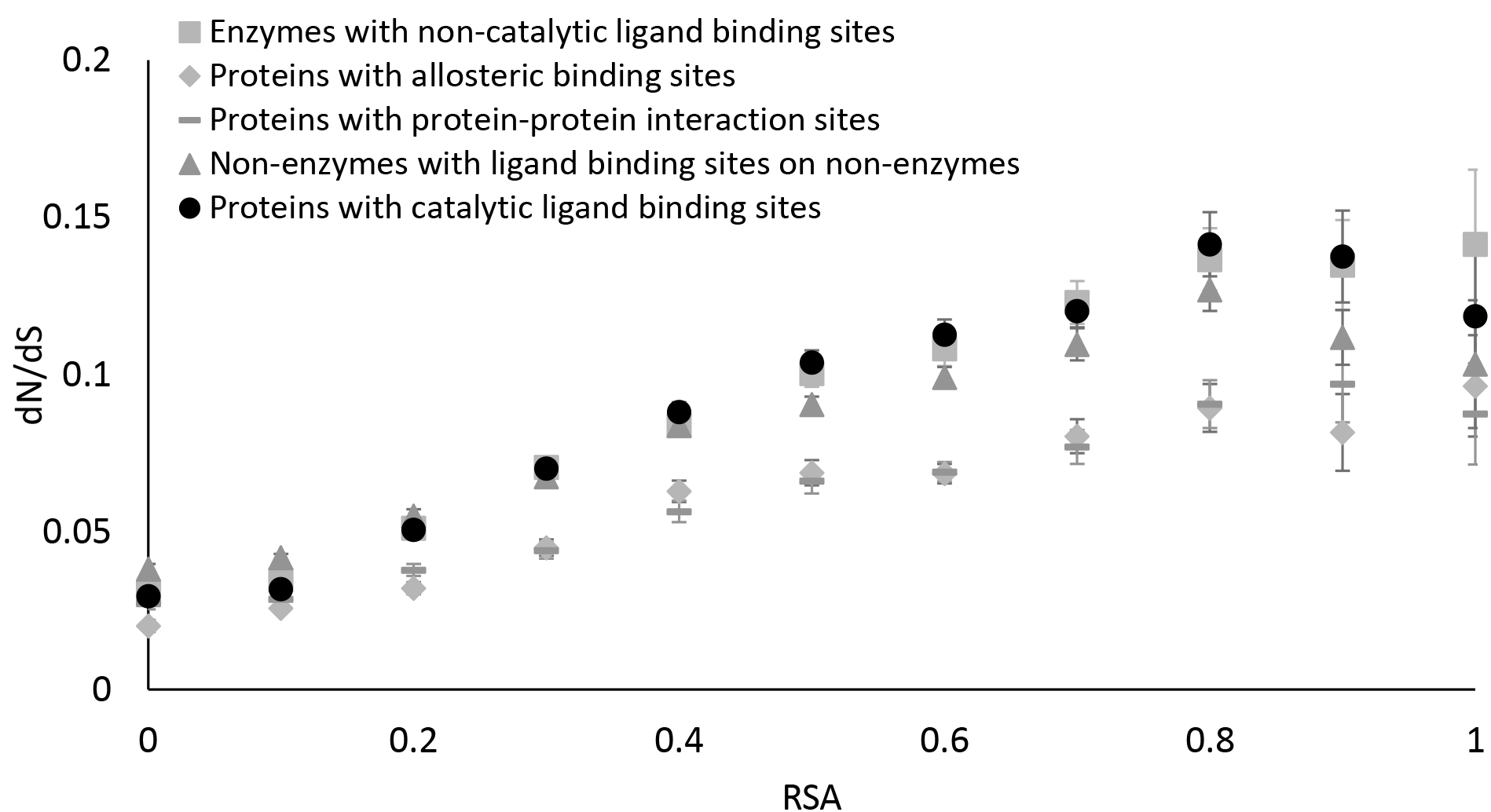
Average evolutionary rate (dN/dS) as a function of relative solvent accessibility (RSA).

**Table S1.** Slope and intercept of the linear fit of the average evolutionary rate (dN/dS) versus RSA.

|  | Slope | Intercept |
| --- | --- | --- |
| Proteins with catalytic binding sites | 0.145 ($\pm 0.011$) | 0.023 ($\pm 0.003$) |
| Enzymes with non-catalytic ligand binding sites | 0.136 ($\pm 0.008$) | 0.026 ($\pm 0.002$) |
| Non-enzymes with ligand binding sites | 0.109 ($\pm 0.006$) | 0.034 ($\pm 0.002$) |
| Proteins with allosteric binding sites | 0.089 ($\pm 0.006$) | 0.018 ($\pm 0.002$) |
| Proteins with protein-protein interaction sites | 0.078 ($\pm 0.004$) | 0.023 ($\pm 0.001$) |

